# A dominant negative mutation uncovers cooperative control of caudal Wolffian Duct development by Sprouty genes

**DOI:** 10.1101/2022.04.07.487514

**Authors:** Gisela Altés, Marta Vaquero, Sara Cuesta, Carlos Anerillas, Anna Macià, Carme Espinet, Joan Ribera, Saverio Bellusci, Ophir D. Klein, Andree Yeramian, Xavi Dolcet, Joaquim Egea, Mario Encinas

## Abstract

The Wolffian Ducts (WD) are paired epithelial tubules central to the development of the mammalian genitourinary tract. Outgrowths from the WD known as the ureteric buds (UB) generate the collecting ducts of the kidney. Later during development, the caudal portion of the WD will form the vas deferens, epididymis and seminal vesicle in males, and will degenerate in females. While the genetic pathways controlling the development of the UB are firmly established, less is known about those governing development of WD portions caudal to the UB. Sprouty proteins are inhibitors of receptor tyrosine kinase (RTK) signaling in vivo. We have recently shown that homozygous mutation of a conserved tyrosine (Tyr53) of Spry1 results in UB defects indistinguishable from that of Spry1 null mice. Here we show that heterozygosity for the Spry1 Y53A allele causes caudal WD developmental defects consisting on ectopically branched seminal vesicles in males and persistent WD in females, without affecting kidney development. Detailed analysis reveals that this phenotype also occurs in Spry1^+/-^ mice but with a much lower penetrance, indicating that removal of tyrosine 53 generates a dominant negative mutation in vivo. Supporting this notion, concomitant deletion of one allele of Spry1 and Spry2 also recapitulates the genital phenotype of Spry1^Y53A/+^ mice with high penetrance. Mechanistically, we show that unlike the effects of Spry1 in kidney development, these caudal WD defects are independent of Ret signaling, but can be completely rescued by lowering the genetic dosage of Fgf10. In conclusion, mutation of tyrosine 53 of Spry1 generates a dominant-negative allele that uncovers fine-tuning of caudal WD development by Sprouty genes.

## INTRODUCTION

Development of the genitourinary system of vertebrates begins shortly after gastrulation through the differentiation of the intermediate mesoderm. This embryonic tissue proliferates, and in some cells mesenchymal-epithelial transition is induced to generate a pair of longitudinal tubules known as the nephric or Wolffian Ducts (WD). Development of the definitive kidney (metanephros) begins on embryonic day (E)10.5 at the caudal end of the WD, where GDNF secreted by the adjacent metanephric mesenchyme prompts the formation of an outgrowth of the WD, the ureteric bud (UB). Subsequently GDNF, through its receptor tyrosine kinase (RTK) Ret, causes the UB to repeatedly branch and ultimately generate the collecting duct system of the kidney. Later during development, the WD will also give rise to the epididymis, vas deferens and seminal vesicle (SV) in males, but will degenerate in females. While the genetic pathways involved in the development of the UB are well-understood, those regulating caudal WD differentiation into male sexual ducts or regression in females are less understood.

The Sprouty family of genes (Spry1-4) encodes feedback inhibitors of RTK activity. In mice, targeted deletion of Spry2 and/or Spry4 causes craniofacial abnormalities due to hypersensitivity to FGF ^1,2^. In addition, genetic deletion of Spry2 leads to hyperplasia of the enteric nervous system as a result of excessive Ret signaling ^3^, whereas Spry1 antagonizes Ret signaling during kidney morphogenesis ^4,5^. Mice deficient in Sprouty1 grow more than one ureteric bud per side of the embryo, owing to excessive activation of Ret signaling ^4,6,7^.

The mechanisms by which Sprouty family members restrain RTK activity are ill-defined. Several in vitro studies have identified a tyrosine located to the N-terminus of the protein that is critical for regulation of Sprouty activity ^8,9,10^. We have recently validated these observations in vivo for the conserved N-terminal tyrosine of Sprouty1, tyrosine 53 ^11^. In that work we show that Spry1 knockin mice lacking tyrosine 53 phenocopy renal defects found in Spry1 knockout animals, consisting on ectopic UB budding and abnormal ureter maturation. The same in vitro studies show that Sprouty mutants lacking this N-terminal tyrosine function as dominant negative alleles. It is however not known whether this behavior is an artifact caused by overexpression.

While characterizing our knockin mice of Sprouty1 lacking tyrosine 53 we observed the emergence of a novel phenotype affecting caudal WD development. Heterozygous Spry1^Y53A/+^ females showed septate vaginas flanked by WDs that did not degenerate but persisted into adulthood, while male mutants exhibited ectopically branched SVs. This phenotype was also observed in Spry1^+/-^ mice of similar genetic background but with much lower incidence, suggesting that mutation of tyrosine 53 generated a dominant-negative allele. Simultaneous removal of one Spry1 and one Spry2 allele recapitulated the Spry1^Y53A/+^ phenotype with high penetrance, indicating that Spry1 and Spry2 collaborate in governing caudal WD development. Finally, we provide in vivo mechanistic data demonstrating that these defects were not caused by excessive Ret signaling as removal of both Ret alleles did not mitigate the phenotype. Instead, removal of one Fgf10 allele completely rescued these abnormalities, pointing to unrestrained signaling by FGF10 as the underlying cause of these defects.

## RESULTS

### Caudal Wolffian Duct defects but normal renal development in Spry1^Y53A^ heterozygous mice

We have recently reported that Sprouty1^Y53ANeo/Y53ANeo^ knockin mice phenocopy renal defects found in Spry1 null mice^11^. When backcrossing Spry1^Y53A/+^ mice into a C57BL/6 genetic background we noticed a sharp decline in fertility of both male and female Spry1^Y53A/+^ mice. Examination of Spry1^Y53A/+^ females in a ∼90% C57BL/6 background revealed that all of them (n=12) had imperforated vaginas and swollen perineum (Figure 1A). Consistently, these females presented hydrometrocolpos, a massively enlarged, fluid-filled uterus occupying most of the abdominal cavity owing to accumulation of uterine gland secretions (Figure 1B). Histological analysis revealed that vaginas from Spry1^Y53A/+^ females presented septa that sometimes resulted in completely duplex vagina (Figure 1C). Moreover, vaginal walls presented paired cystic cavities running parallel to vaginal lumens, lined by monostratified epithelium, similar to WD remnants known as Gartner cysts in humans (Figure 1C). On the other hand, Spry1^Y53A/+^ males in the same genetic background presented normal testes, epididymes and vasa deferentia (data not shown), but SVs were duplicated or presented ectopic branches in most animals (Figure 1D). Microscopic examination showed normal histological organization of the SV (Figure 1D). Of note, gross morphology of kidneys and ureters appeared normal in these mice (Figure 1B).

**Figure 1.**
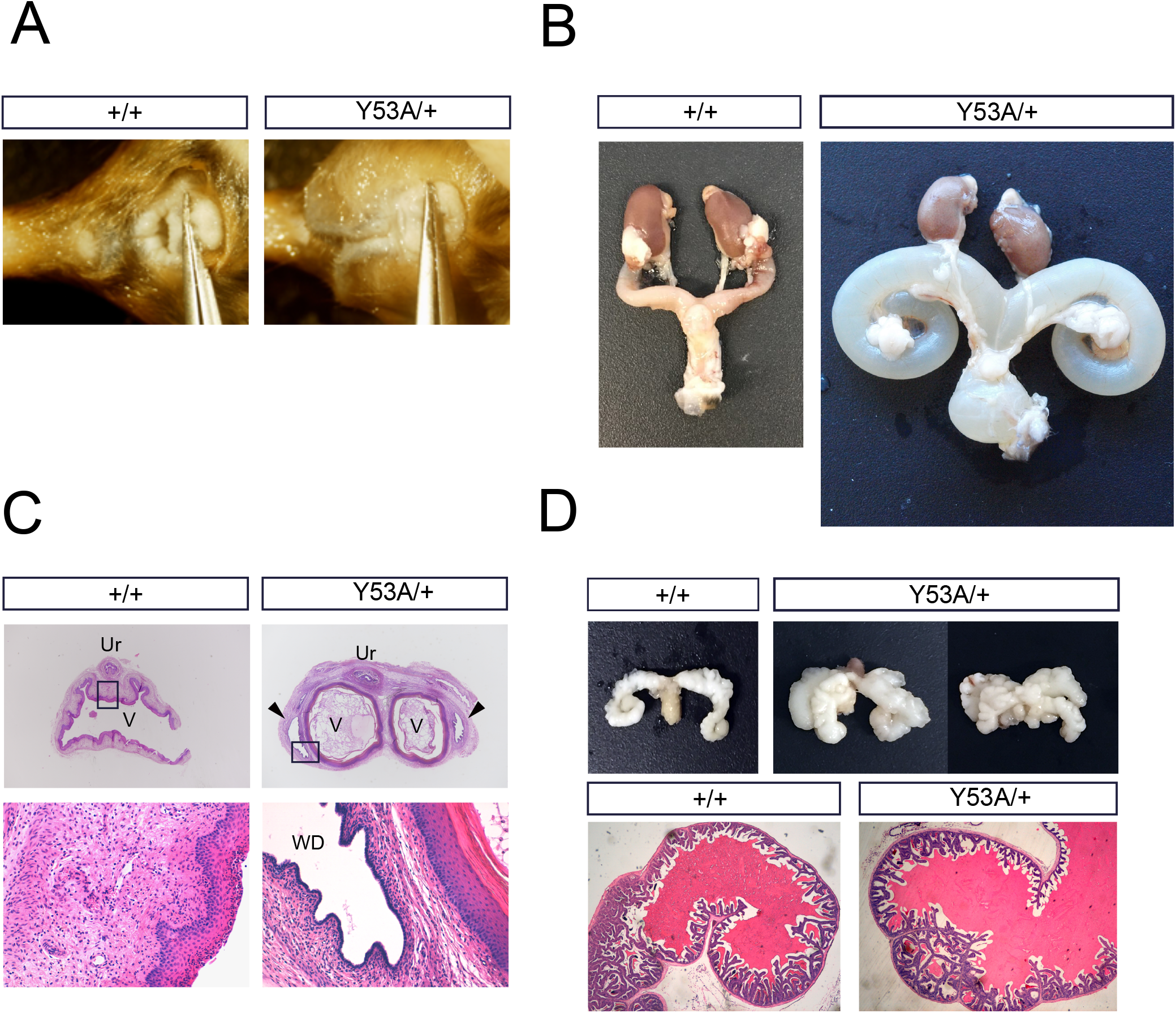
Mutation of Spry1 tyrosine 53 perturbs caudal WD development. Adult Spry1^Y53A/+^ mice have imperforate vagina (A) and hydrometrocolpos (B) but normal ureters and kidneys. (C) Hematoxylin-eosin stained paraffin sections of vaginas from adult females of the indicated genotypes. Mutant females show septate vaginas flanked by tubular cavities lined with a monostratified epithelium, reminiscent of human Gartner cysts (arrowheads). Insets show higher magnification pictures. (D) SV from mutant mice present ectopic branches but normal histology in adult Spry1^Y53A/+^ mice. Ur, urethra; V, vagina.

We next performed whole-mount cytokeratin staining at different key embryonic ages to characterize development of the genitourinary system of these mice. No differences in UB budding or branching were observed between wild type and Spry1^Y53A/+^ animals at E11.5 and E13.5, respectively (Figure 2A, top and middle panels). Ureter maturation was also normal in Spry1^Y53A/+^ mice (Figure 2A, bottom panel, and Supplemental Videos 1 and 2). As we previously described^11^, only a small proportion of heterozygous animals displayed single ectopic UBs leading to unilateral duplex ureters that unlike those from Spry1^Y53A/Y53A^ mice correctly separated from the WD (Supplemental Figure 1A, B). The presence of WD remnants in adult Spry1^Y53A/+^ females prompted us to investigate whether WD degeneration failed along the whole length of the tubules or just at their caudal-most portion. As shown in Figure 2B, WD degenerated normally throughout most of their length, except for the portions located at the level of the vagina. These remnants were clearly present at birth in heterozygous animals but absent in wild type littermates (Figure 2B and Supplemental Video 3). In males, WD maturation appeared roughly normal up to E16.5, but showed ectopic branching at birth (Figure 2C). In conclusion, abnormal caudal WD development appeared in late stages of embryonic development of Spry1^Y53A/+^ mice, which display largely normal urinary tract development.

**Figure 2.**
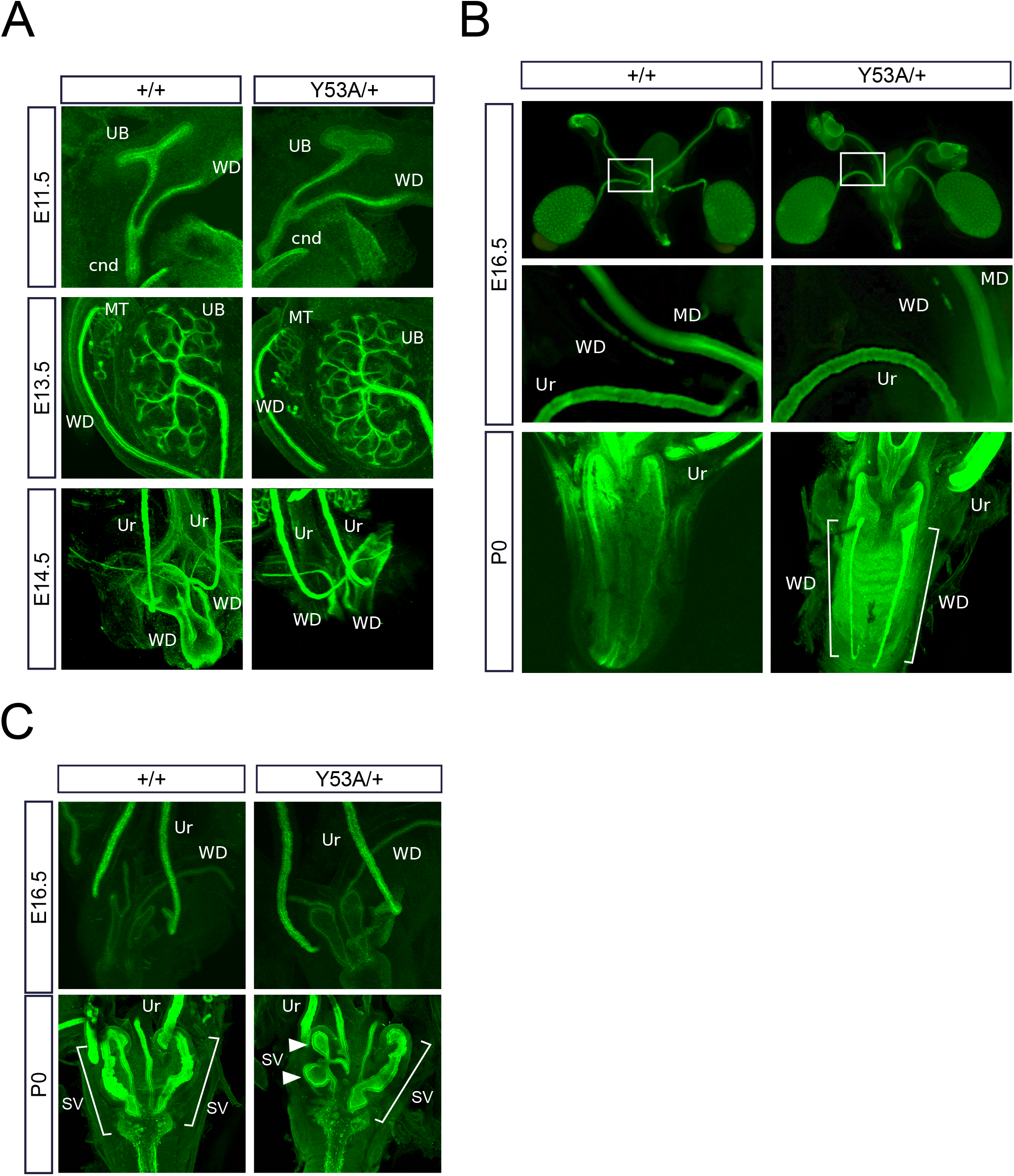
Developmental defects of WD-derived structures of Spry1^Y53A/+^ mice are restricted to its caudal-most portion. Cytokeratin staining of genitourinary systems of Spry1^Y53A/+^ embryos of the indicated ages reveals (A) normal UB budding, branching and ureter maturation, (B) proper degeneration of cranial WD (top two panels) and abnormal retention of the WD at the level of the vagina (bottom panels), and (C) aberrant SV branching at birth (arrowheads). Cnd, common nephric duct; MD, Müllerian Duct; MT, mesonephric tubules; Ur, ureter.

### Mutation of Spry1 tyrosine 53 generates a dominant-negative allele

Previous data demonstrate that overexpressed Sprouty mutants lacking their N-terminal tyrosine act as dominant negative mutants in vitro^8–10^. To ascertain if the same was true in vivo, we compared the penetrance of the above caudal WD defects in Spry1^Y53A/+^ vs. Spry1^+/-^ mice. To control for genetic background effects, we analyzed both genotypes in ∼90% as well as ∼40% mixed C57BL/6×129Sv background (see Supplemental Figure 1 for details on the crosses performed). As shown in Figure 3A, the caudal WD phenotype was not restricted to Spry1^Y53A/+^ animals but was also found in Spry1^+/-^ mice. Second, in both Spry1^Y53A/+^ and Spry1^+/-^ mice the phenotype was dependent on the genetic background, being more penetrant in purer C57BL/6 backgrounds. Finally, the penetrance of caudal WD abnormalities was always higher in Spry1^Y53A/+^ than in Spry1^+/-^ mice of similar genetic background, thus meeting the definition of a dominant negative allele.

**Figure 3.**
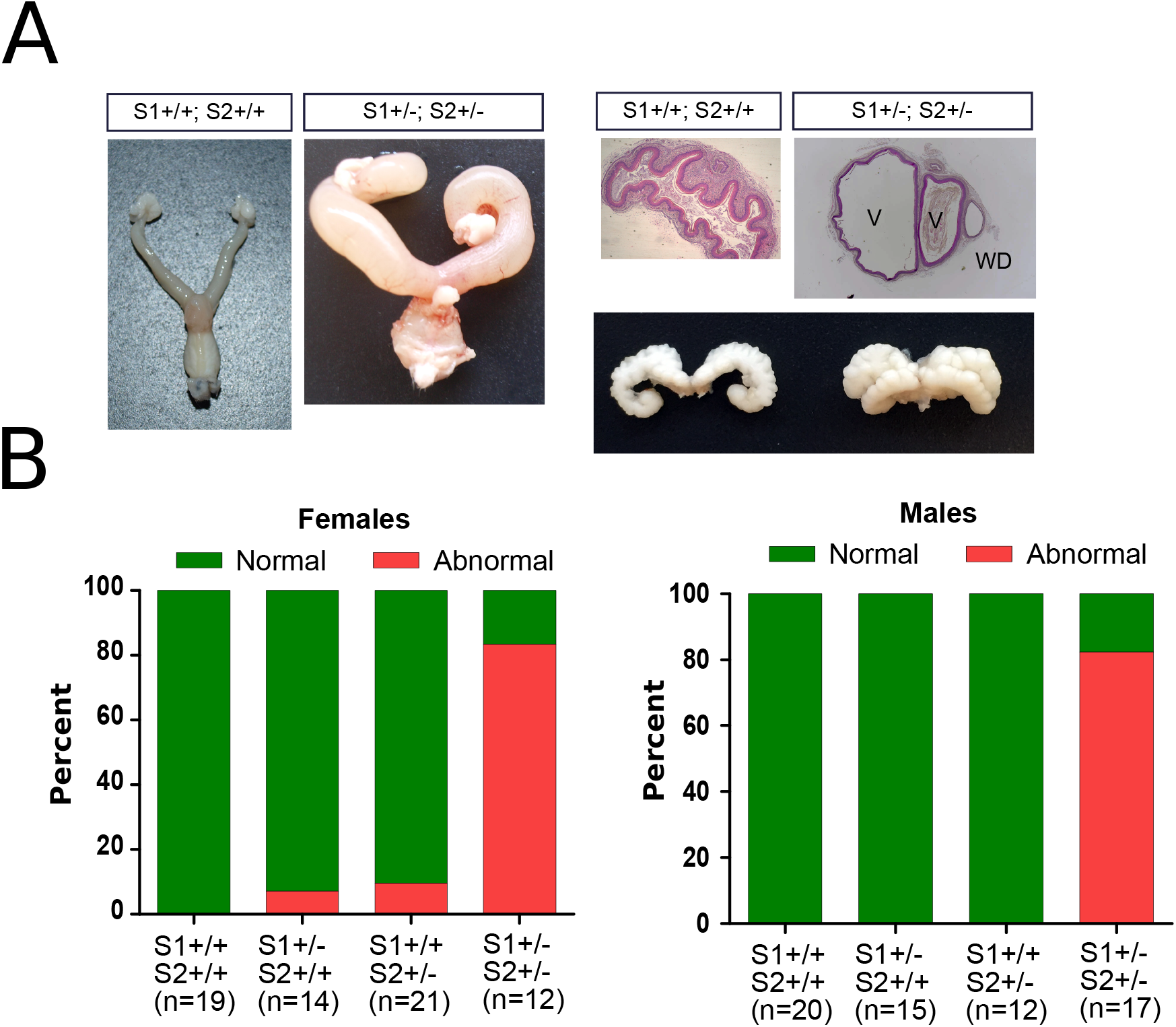
Mutation of Spry1 tyrosine 53 generates a dominant negative allele. (A) Frequencies of the caudal WD defects found in adult mice of the indicated genotypes and genetic background. Note that for a given genetic background the penetrance of the defects is always higher in Spry1^Y53A/+^ than in Spry1^+/-^ mice, indicating a dominant negative behavior. (B) Spry1 protein levels are higher in SV from Spry1^Y53A/+^ (n=7) newborn animals than in wild type (n=4) littermates as assessed by immunoblot. (C) Left panel, densitometric analysis of the samples above shows a roughly three-fold increase of levels in mutant animals. Right panel, mRNA levels of Spry1 in SV from newborn mice are not significantly different between Spry1^Y53A/+^ (n=19) and Spry1^+/+^ (n=12) mice. In both cases, indicated p-values were calculated using Mann-Whitney’ s U test.

One of the mechanisms by which loss-of-function mutants behave as dominant negatives is that their expression levels are abnormally high, thus interfering with normal functioning of signaling pathways (for example competing with wild type molecules for upstream activators). We therefore examined Spry1 protein levels in the SV of Spry1^Y53A/+^ mice vs Spry1^+/+^ littermates, and found much higher levels in the former (Figures 3B and 3C). Such an increase was post-transcriptional since mRNA levels were comparable between phenotypes (Figure 3C). This is in agreement with a robust increase of Spry1 protein (but not mRNA) levels in the developing kidney of Spry1^Y53ANeo/Y53ANeo^ mice ^11^. Thus, Spry1 Y53A could exert its dominant negative effects at least partially via elevated protein levels.

### Spry1 and Spry2 cooperate to pattern internal genitalia

Spry1 and Spry2 have overlapping functions during development of a variety of structures such as the cerebellum^12^, inner ear^13^, eyelids^14^, teeth^15^, or external genitalia^16^, among others. To explore whether Spry1 and Spry2 also collaborate in patterning of the caudal WD, we crossed Spry1^+/-^ mice to Spry2^+/-^ mice to obtain double heterozygous animals. Examination of external genitalia of twelve double heterozygous females revealed that ∼80% of them had imperforated vagina with hydrometrocolpos, and septate vaginas flanked by WD remnants (Figure 4), despite showing correctly formed kidneys and lower urinary tracts (not shown). Likewise, double heterozygous males presented duplicated SVs in a similar proportion (∼80%, n=17; Figure 4), with no other apparent abnormalities in testis, epididymis and vas deferens. Importantly, the penetrance of such caudal WD defects in single heterozygous mice was negligible, according to its mixed genetic background (50% C57BL/6×129Sv). Thus, removal of one allele each of Spry1 and Spry2 resulted in a large percentage of animals displaying a caudal WD phenotype identical to that observed in Spry1^Y53A/+^ mice, demonstrating that Spry1 and Spry2 collaborate with each other during development of the caudal WD.

**Figure 4.**
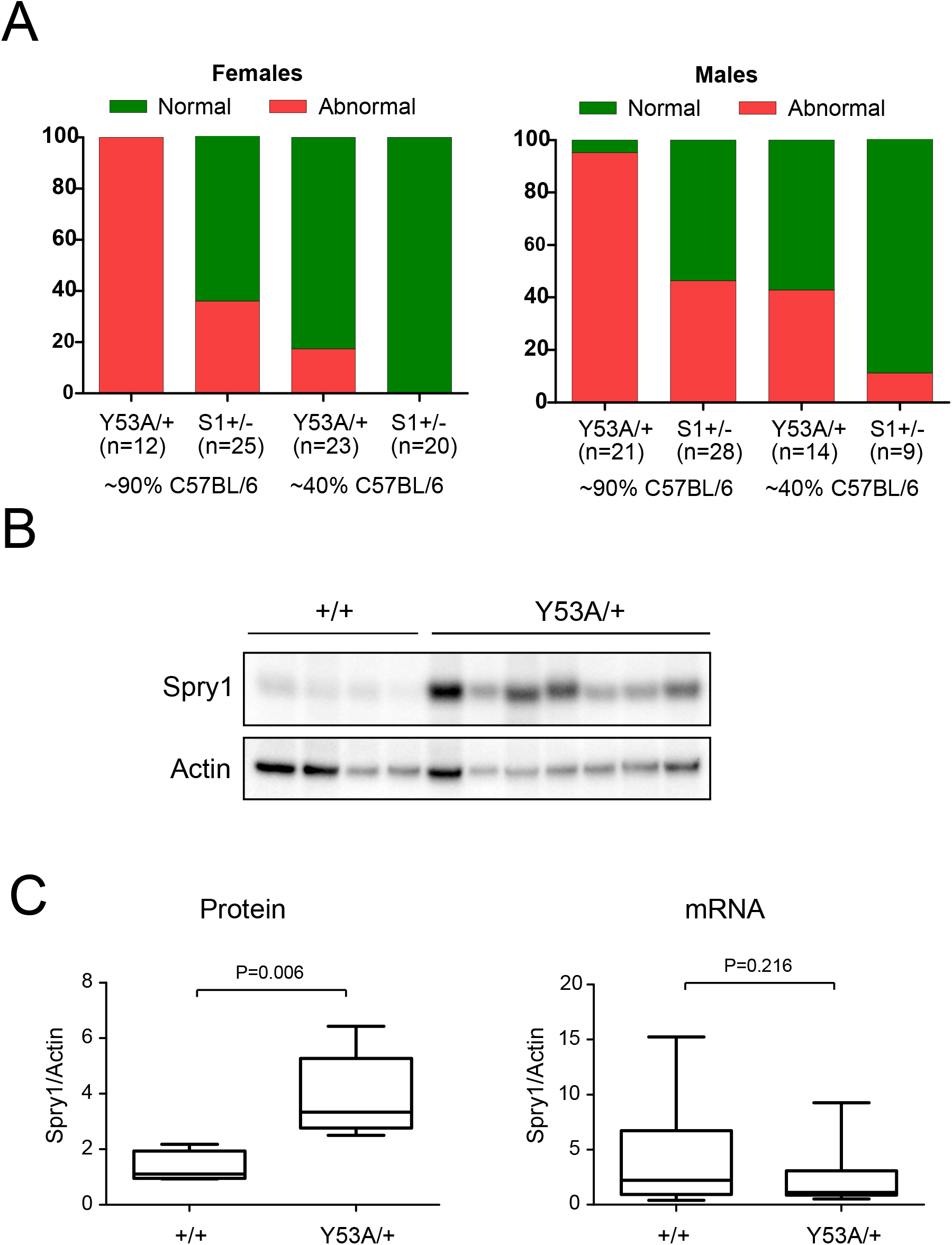
Spry1 and Spry2 cooperate to determine caudal WD fate. (A) Spry1, Spry2 double heterozygous mice present defects identical to those found in Spry1^Y53A/+^ including blind, septate vagina flanked by WD remnants and duplex seminal vesicle. (B) Phenotypic frequencies of caudal WD defects in adult animals of the indicated sex and genotype. Double heterozygous mice present a much higher penetrance of these defects than single heterozygous mice.

### Caudal Wolffian duct defects are Ret-independent

Since Spry1 antagonizes Ret signaling during ureteric bud outgrowth and branching, we wanted to analyze whether upregulated Ret activity underlies these caudal WD defects. We first examined the expression pattern of Ret in the caudal WD during development. To do so, we took advantage of knockin mice expressing EGFP from the Ret locus (Ret^EGFP/+^, Figure 5A)^17^. In males, EGFP fluorescence was localized to the caudal WD from E14.5 to E16.5, and persisted in SVs from birth to at least the first postnatal week (Figure 5B). In wild type females the WD was positive for EGFP fluorescence until it regressed at around E16.5 (Figure 5C, left panel). We then generated Spry1^+/-^; Spry2^+/-^; Ret^EGFP/+^ triple mutant mice and analyzed their caudal WD remnants by EGFP fluorescence at birth. As shown in Figure 5C (right panel), WD remnants found in newborn mutant females were also positive for Ret.

**Figure 5.**
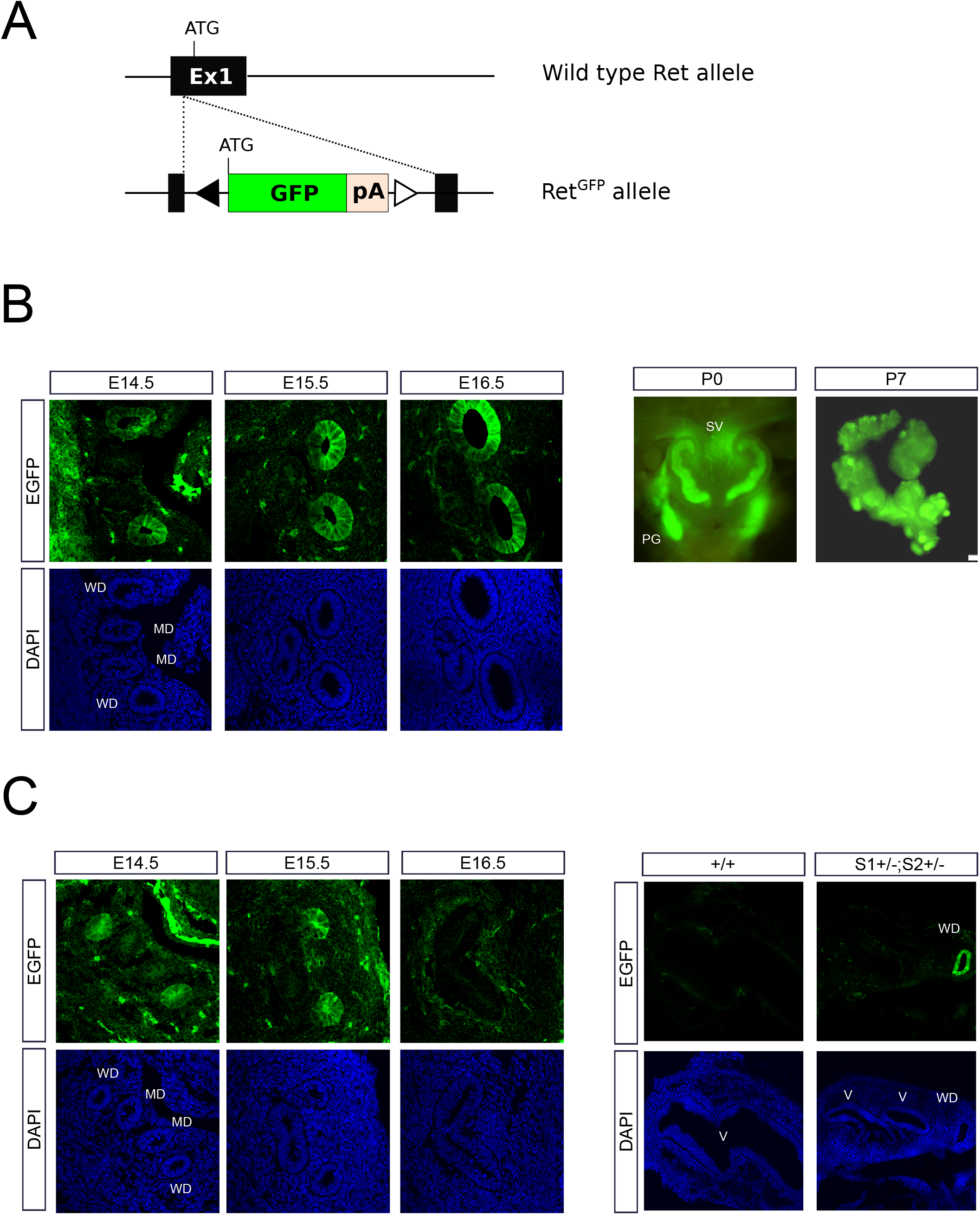
Ret is expressed in the caudal-most portions of the WD. (A) Diagram showing the structure of the Ret^EGFP^ allele used to detect Ret expression. Note that expression of endogenous Ret from this allele is ablated. (B) Ret is expressed in caudal WD from male embryos throughout development and postnatally in SV. (C) Ret is expressed in degenerating WD in wild type females (left panel), and in WD remnants of Spry1/2 double heterozygous newborn mice (right panel). MD, Müllerian Duct; PG, pelvic ganglia (express Ret); SV, seminal vesicle; V, vagina.

We then generated Spry1^+/-^; Spry2^+/-^ mice either expressing Ret (Ret^EGFP/+^) or not (Ret^EGFP/EGFP^) and analyzed their caudal WDs by EGFP fluorescence at birth, since Ret knockout animals die shortly after birth owing to renal agenesis (Figure 6). In pups wild type for Spry1 and Spry2 only one SV per side of the embryo was formed, irrespective of the Ret genotype (Ret^EGFP/+^ or Ret^EGFP/EGFP^; Figure 6A), indicating that Ret is dispensable for patterning of the SV. More importantly, caudal WD defects found in Spry1 and Spry2 double heterozygous mice were unaffected by deletion of Ret in both male and females pups (Figure 6A and B). Taken together, these observations indicate that caudal WD defects responsible for the internal genitalia abnormalities presented by Sprouty mutant mice are not caused by dysregulated Ret signaling. This is in sharp contrast to what happens with UB development, which is very sensitive to changes in the GDNF-Ret signaling axis.

**Figure 6.**
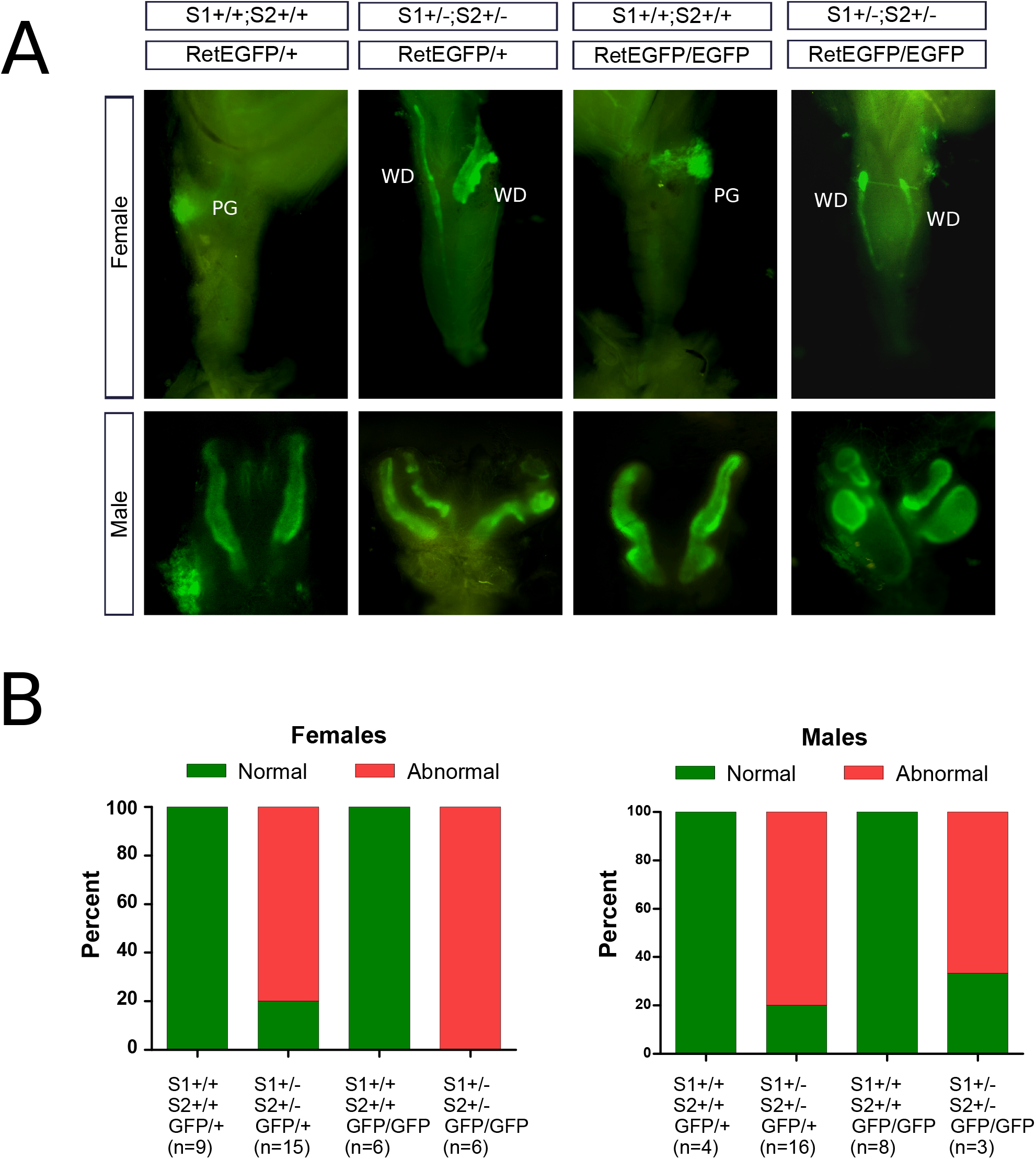
Caudal WD defects are Ret-independent. (A) EGFP fluorescence of vaginas and seminal vesicles form newborn mice of the indicated genotypes. Quantification is shown in the lower panel (B). Note normally formed vaginas and SV in animals expressing wild type Spry1 and Spry2 irrespective of the Ret genotype. In Spry1^+/-^; Spry2^+/-^ mice, WD remnants and duplex seminal vesicles are not rescued by loss of Ret expression.

### Caudal WD defects are completely rescued by heterozygous deletion of Ffg10

Since aberrant Ret signaling did not appear to contribute to caudal WD defects, we sought to explore whether dysregulated activity of other RTKs could be responsible for them. To narrow down putative candidates, we performed RNAseq on pooled SVs from wild type newborn animals and examined RTK expression. As shown in Supplemental Table 1, Fgfr2 and Fgfr1 were among the top 5% most abundant genes in neonatal SV, ranking as the third and fifth most expressed RTKs respectively. Moreover, FGF7 and FGF10 have been shown to promote branching morphogenesis in organ culture of SVs^18,19^, and WD retention in explanted female metanephroi^20^. We analyzed whether our phenotype could be rescued by attenuating Fgf10 signaling since its expression roughly doubled that of Fgf7 in neonatal SVs (Supplemental Table 2). To do so, we generated either Spry1^Y53A/+^ or Spry1^+/-^; Spry2^+/-^ mice lacking one copy of Ffg10. The former were generated by crossing Spry1^Y53A/+^ mice to Fgf10^+/-^ mice, whereas the latter were produced by mating Spry1^f/f^; Spry2^f/f^; UbC-Cre^ERT2^ male mice injected with tamoxifen at four weeks of age to Fgf10^+/-^ females (therefore, only Spry1^+/-^; Spry2^+/-^; Fgf10^+/+^ or Spry1^+/-^; Spry2^+/-^; Fgf10^+/-^ were generated, see Supplementary Figure 2). As shown in Figure 7, genetic ablation of a single copy of Fgf10 completely rescued the caudal WD phenotype caused by both heterozygous expression of Spry1Y53A and double heterozygous deletion of Spry1 and Spry2, indicating that loss of function of Spry genes cause the caudal WD phenotype at least in part by exacerbating FGF10-mediated downstream signaling.

**Figure 7.**
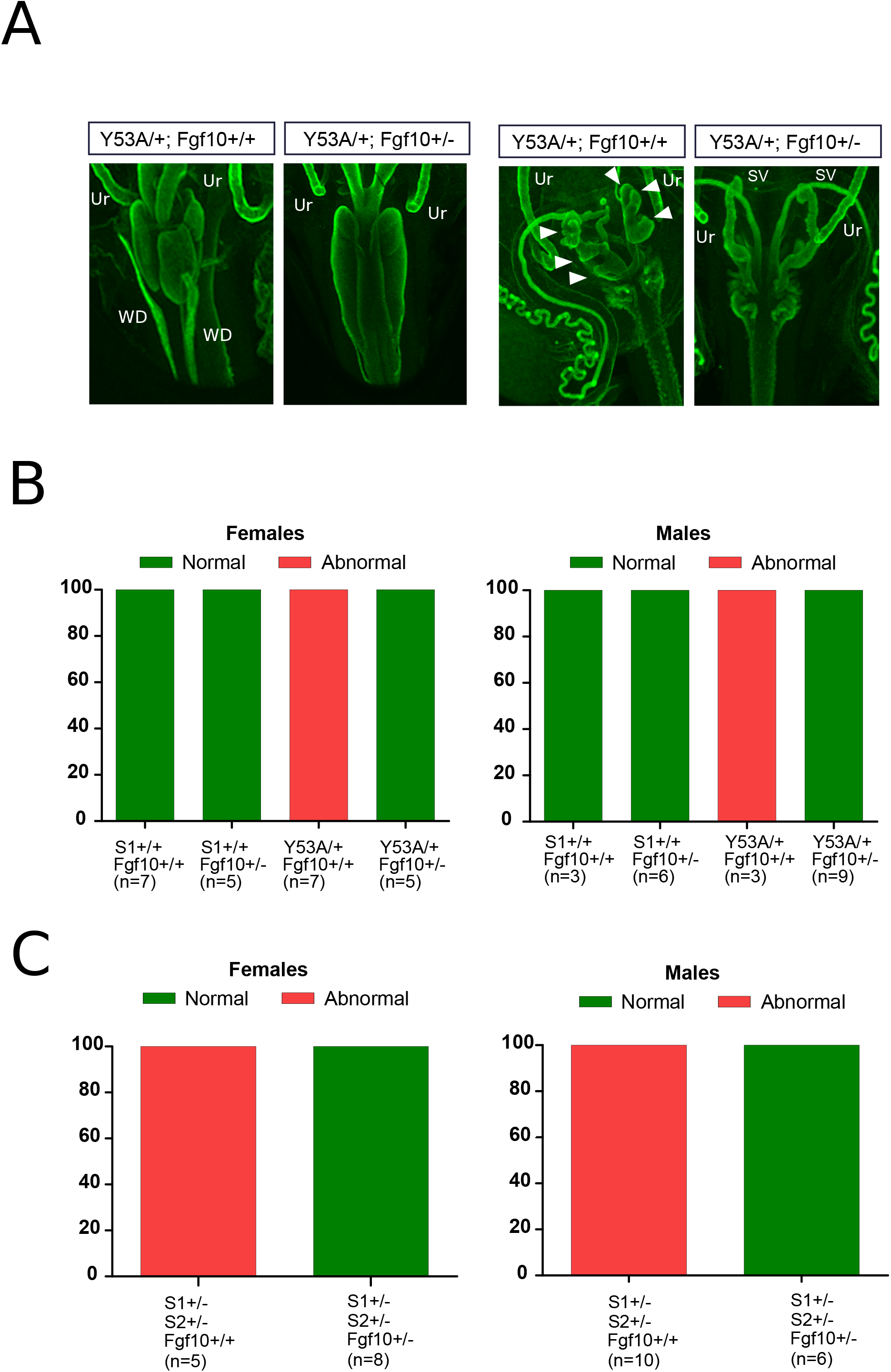
Heterozygous deletion of Fgf10 completely rescues WD defects of Spry1^Y53A/+^ and Spry1^+/-^; Spry2^+/-^ mice. (A) Whole mount cytokeratin staining of female (two left panels) and male (two right panels) newborn mice of the indicated genotypes. Arrowheads point to ectopic SV branches. Ur, ureter. (B, C) Frequencies of caudal WD defects of newborn mice of the indicated genotypes reveal total phenotype rescue upon heterozygous deletion of Fgf10.

## DISCUSSION

In this work we report a novel function for Sprouty proteins in patterning internal genitalia by controlling caudal WD development. This previously unnoticed role was revealed by the dominant-negative nature of Spry1Y53A mutation, which in contrast to the null mutation, causes the emergence of the phenotype with high penetrance. We also report that removing one allele of Spry2 in the context of Spry1 heterozygosity recapitulates this WD phenotype with high penetrance. Unlike renal malformations, these WD abnormalities are independent of Ret signaling but are rescued by decreasing the genetic dosage of Fgf10.

Removal of the N-terminal tyrosine of Sprouty proteins has been shown to generate dominant negative mutants in vitro. Thus Sasaki et al ^8^ reported that expression of either Y53A Spry4 or Y55A Spry2 reverses the inhibitory effect of wild type Spry2 or Spry4 on ERK activation, respectively. Furthermore, expression of HA-tagged Y55F Spry2 blocks FGF-induced tyrosine phosphorylation of myc-tagged Spry2 or Spry1 in a dose-dependent manner ^21^. We have found that Spry1Y53A protein levels are much higher than those of wild type protein in SVs, thus providing a simple mechanism by which the mutant behaves as a dominant negative.

Why are protein levels of Spry1 Y53A higher? Compelling evidence show that binding of c-Cbl to the N-terminal tyrosine of human Spry2 promotes its ubiquitin-dependent degradation ^22,23,24^. We speculate that analogously, lack of Tyr53 of Spry1 results on its accumulation owing to defective binding to c-Cbl and compromised proteasomal clearance. Higher steady-state levels of mutant Spry1 protein would lead to dominant negative effects, caused by saturation of upstream regulators and/or downstream effectors by the inactive protein.

We have found that Spry1 and Spry2 collaborate in governing caudal WD development. While overlapping activities of Spry1 and Spry2 have been frequently described, others appear to be specific for either protein. For example, Spry1 specifically controls ureteric bud formation ^4^ whereas Spry2 regulates enteric nervous system or inner ear development ^3;25^. Since the structure of both proteins is highly similar, and given that Sprouty proteins have been previously shown to hetero-oligomerize ^10,21^, one likely explanation for these observations is that expression levels in different tissues rather than distinct mechanisms of action would determine the importance of each member in a given tissue (noteworthy, oligomerization depends on the C-terminal, cysteine-rich domain but not on the N-terminal tyrosine^10,21^). Based on all the above data, together with the observation that Spry1 behaves as a dosage-sensitive gene^6,26^, we propose a model in which global output of Spry1/2 activity would dictate the fate of WD-derived structures (Figure 8).

**Figure 8.**
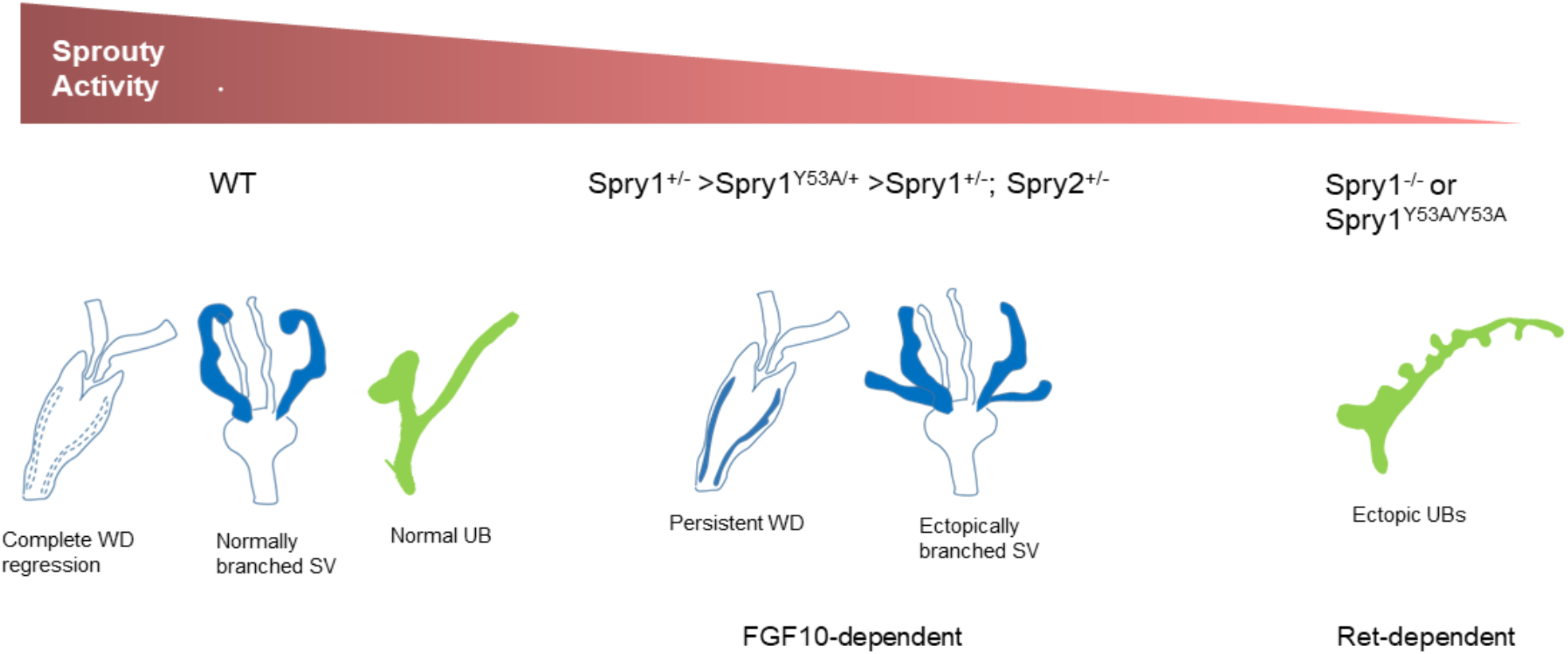
Global Spry1/2 activity and tissue sensitivity govern development of the WD. We propose a model in which global, quantitative Spry1/2 activity and tissue sensitivity towards this activity determines the fate of WD-derived structures. Wild type animals show full activity whereas Spry1^+/-^, Spry1^Y53A/+^, Spry1^+/-^; Spry2^+/-^ and Spry1^-/-^ or Spry1^Y53A/Y53A^ display progressively decreasing Spry activity. SV and WD remnants are relatively sensitive to diminished activity, whereas UB requires very low activity to disrupt its normal development. On the other hand, caudal WD development relies on FGF10 signaling whereas UB development is mainly dependent on Ret activity.

Caudal WD abnormalities were not rescued by removing both alleles of Ret, indicating that unlike UB defects, they were not caused by unrestrained Ret signaling. Instead, removing one single copy of Fgf10 completely prevented their emergence. Our RNAseq data show that Fgfr2 and to a lesser extent Fgfr1 are abundantly expressed in the mouse neonatal SV. In situ hybridization experiments show that Fgfr2 is robustly expressed in the caudal WD epithelium but not its surrounding mesenchyme whereas Fgfr1 is expressed in the mesonephric tubules and the caudal WD mesenchyme during development^27,28^. On the other hand, signaling by Fgfr2 supports caudal WD maintenance, as conditional deletion of Fgfr2 in the WD epithelium using HoxB7-Cre transgenic mice results in regression of the caudal, but not cranial WD^28^. Interestingly, the UB appeared to develop normally in these mice. In line with these pieces of evidence, Kuslak and colleagues^27^ have demonstrated that the spontaneous mouse SV shape (svs) mutation, which affects budding/branching of the SV is caused by a mutation of the Fgfr2 gene that perturbs splicing of the gene. Authors also show that the alternative splice isoform Fgfr2c, whose cognate ligands are FGF7 and FGF10 ^29^, is the main isoform expressed in the SV^27^. Previous work has shown that both FGF7 (also known as KGF) and FGF10 are expressed in the mesenchyme surrounding SVs during development and, as mentioned earlier, that both factors can promote SV branching in vitro^18,19^. Likewise, exogenous FGF7 and/or FGF10 causes retention of the WD in explanted female mesonephroi at different embryonic ages^20^. These data suggest that both FGF7 and FGF10 play a role controlling caudal WD development. However, while the phenotype of FGF7 knockout mice is relatively mild with no reported defects in internal genitalia^30^, deletion of FGF10 dramatically affects both prostate and SV development^31^. In light of these data and our own observations, we conclude that FGF10 is the major regulator of caudal WD development, although a redundant role of FGF7 cannot be excluded.

What are the cellular mechanisms controlling WD regression in females? Two key reports demonstrate that several structures in the developing embryo including the WD in females degenerate via induction of (programmed) cellular senescence ^32,33^. This type of cellular senescence relies on expression of the cdk inhibitor p21, and its transcriptional signature is characterized by increased activation of the TGFβ, Shh and Wnt/β-catenin pathways ^32^. Interestingly, adult female mice lacking p21 show septate vagina, although persistent WD was not examined^34^. Interestingly, the phenotype of our Spry mutants is strikingly similar to that of a recently described β-catenin mutant, both in males and females^35^. An important next step will be investigating the relationship between Sprouty proteins, the Wnt/β-catenin pathway and cellular senescence during development of the caudal WD.

## METHODS

### Mice

All animal use was approved by the Animal Care Committee of the University of Lleida in accordance with the national and regional guidelines. Mice were maintained on a 12h light/dark cycle, and food and water was provided ad libitum. Spry1 Y53A (Spry1^tm1.1Mns^), Spry1 floxed (Spry1^tm1Jdli^), and Spry2 floxed (Spry2^tm1Mrt^) mice have been previously described ^4,11,25,36^. Fgf10 null mice were generated^37^ by crossing Fgf10 floxed mice (Fgf10^tm1.2Sms/J^) to CMV-Cre mice [B6.C-Tg(CMV-cre)1Cgn/Jas]. Spry2 knockout mice [Spry2^tm1.1Mrt, 25^] were obtained from the Mutant Mouse Resource & Research Center (MMRRC, https://www.mmrrc.org). Ubiquitin C-Cre^ER^ transgenic mice (Ndor1^Tg(UBC-cre/ERT2)1Ejb^) were from the Jackson Laboratories (Bar Harbor, ME, USA). Spry1 knockout (Spry1^tm1.1Jdli, 4^) and Ret EGFP mice (Ret^tm13.1Jmi^,^17^) were generous gifts from Dr. Albert M. Basson (King’ s College, London, UK), and from Dr. Sanjay Jain (Washington University, St Louis, USA), respectively.

### Whole mount staining

For EGFP fluorescence, specimens were dissected and directly photographed under a Nikon SMZ18 fluorescence stereoscope coupled to a Nikon DS-Ri2 camera. For whole mount cytokeratin staining, tissues were fixed overnight at 4ºC in 4% paraformaldehyde, blocked in blocking buffer (4% BSA, 1% Triton X100,100 mM glycine, 0.2% sodium azide in PBS) overnight at 4ºC with gentle agitation and incubated with a 1:100 dilution of anti-cytokeratin antibody (TROMA-I, DSHB) in blocking buffer for 3-5 days at 4ºC. In some experiments, retrieval using FLASH reagent 2 was performed before blocking as described^38^. In these experiments, blocking buffer was 10% FBS, 1% BSA and 5% DMSO in PBS containing 0.2% Triton X100. After primary antibody incubation, specimens were washed three times in PBS 1% Triton X100 for 2 hours each at room temperature, and incubated with fluorescently labeled secondary antibodies (Jackson Immunoresearch) diluted 1: 100 in blocking buffer overnight at 4ºC. Next day, tissues were washed again three times in PBS 1% Triton X100 for 2 hours at room temperature. Tissues were cleared in 1:2 benzyl alcohol:benzyl benzoate (BABB) after being dehydrated through a graded series of 25%, 50%, 75% and 100% methanol in PBS (1h at room temperature each). Specimens were placed on coverslips and imaged using a Olympus Fluoview FV1000 confocal laser scanning microscope or a Nikon SMZ18 fluorescence stereoscope.

### Western blot

Protein extracts from tissues were obtained by mechanical lysis using denaturing lysis buffer (50 mM HEPES, 2% SDS), boiled and sonicated. Protein was electrophoresed on 10% SDS polyacrylamide gels, transferred onto PVDF membrane (Millipore) and blocked using 3% BSA (Sigma) in TBS-Tween (0.1%, TBST) for 1h at room temperature. Membranes were incubated with primary antibodies overnight at 4ºC. Signal was detected using Horseradish Peroxidase (HRP)-conjugated secondary antibodies (1:10000, Jackson ImmunoResearch), followed by a chemiluminescence reaction using Amersham ECL substrate (GE Healthcare). Chemiluminescent signal was detected using the VersaDoc Imaging system Model 4000 (Bio-Rad), and densitometry was performed using its software package (ImageLab, Bio-Rad). Primary antibodies were Anti-Spry1 (Cell Signalling, Cat# 13013 or Cat# 12993) and Anti β-actin (Santa Cruz, sc-1616).

### RT-qPCR

Total RNA was extracted with TRIZOL reagent (ThermoFisher), by homogenizing snap frozen tissue using a TissueLyser LT device (Qiagen). RNA was reverse transcribed using the High-Capacity cDNA Reverse Transcription Kit (ThermoFisher) as per manufacturer’ s instructions. Quantitative RT-PCR (RT-qPCR) reactions were performed by means of the SYBR green method, using either the 2x Master mix qPCR Low Rox kit (PCR Biosystems). The 2^-ΔΔCt^ method was used, normalizing to actin expression. Reverse transcriptase-minus and blank reactions were included in all experiments. Primers used were as follows: Spry1 Fwd 5’ -CTCTGCGGGCTAAGGAGC-3’ ; Spry1 Rev 5’ - ACGCCGGCTGATCTTGC-3’ ; Actin Fwd 5’ - TTCTTTGCAGCTCCTTCGTT-3’ ; Actin Rev 5’ -ATGGAGGGGAATACAGCCC-3’.

## ACKNOWLEDGEMENTS

We are grateful to Dr. Albert M. Basson (King’ s College, London), Dr. Sanjay Jain (Washington University, St Louis) and Dr. Daniel Sanchis (Universitat de Lleida) for sharing valuable reagents. We thank Marta Hereu, for her excellent technical assistance. This work was supported by grants BFU2017-83646-P (MINECO) and PID2020-114947GB-I00 (MCIU) (both supported by funds from AEI/FEDER, UE) to ME. MV was supported by a predoctoral fellowship from AGAUR. GA and CA and GA are supported by a fellowship from Universitat de Lleida. SC was supported by a Cofund action from the Marie Curie program of the EU.

## AUTHOR CONTRIBUTIONS

Conceptualization, GA, MV, ME; Methodology, GA, MV, CE, JR, SB, ODK, XD, JE, ME; Investigation, GA, MV, SC, CA, AM, CE, JR, AY, XD, JR, ME; Writing–original draft, ME; Writing–Review & Editing, MV, GA, CE; JR, SB, ODK, XD, JE, ME Visualization, MV, GA, ME; Supervision, ME; Funding Acquisition, ME.

## REFERENCES

1. Masoumi-Moghaddam, S., Amini, A. & Morris, D. L. The developing story of Sprouty and cancer. Cancer metastasis reviews 33, 695–720 (2014).

2. Neben, C. L., Lo, M., Jura, N. & Klein, O. D. Feedback regulation of RTK signaling in development. Developmental Biology (2017) doi:10.1016/j.ydbio.2017.10.017.

3. Taketomi, T. et al. Loss of mammalian Sprouty2 leads to enteric neuronal hyperplasia and esophageal achalasia. Nat Neurosci 8, 855–857 (2005).

4. Basson, M. A. et al. Sprouty1 is a critical regulator of GDNF/RET-mediated kidney induction. Dev Cell 8, 229–239 (2005).

5. Basson, M. A. et al. Branching morphogenesis of the ureteric epithelium during kidney development is coordinated by the opposing functions of GDNF and Sprouty1. Dev Biol 299, 466–477 (2006).

6. Rozen, E. J. et al. Loss of Sprouty1 rescues renal agenesis caused by ret mutation. Journal of the American Society of Nephrology 20, (2009).

7. Michos, O. et al. Kidney development in the absence of Gdnf and Spry1 requires Fgf10. PLoS Genet 6, e1000809 (2010).

8. Sasaki, A., Taketomi, T., Wakioka, T., Kato, R. & Yoshimura, A. Identification of a dominant negative mutant of Sprouty that potentiates fibroblast growth factor-but not epidermal growth factor-induced ERK activation. J Biol Chem 276, 36804–36808 (2001).

9. Hanafusa, H., Torii, S., Yasunaga, T. & Nishida, E. Sprouty1 and Sprouty2 provide a control mechanism for the Ras/MAPK signalling pathway. Nat Cell Biol 4, 850–858 (2002).

10. Mason, J. M. et al. Tyrosine Phosphorylation of Sprouty Proteins Regulates Their Ability to Inhibit Growth Factor Signaling: A Dual Feedback Loop. Molecular Biology of the Cell 15, 2176–2188 (2004).

11. Vaquero, M. et al. Sprouty1 Controls Genitourinary Development via its N-Terminal Tyrosine. Journal of the American Society of Nephrology 30, 1398–1411 (2019).

12. Yu, T., Yaguchi, Y., Echevarria, D., Martinez, S. & Basson, M. A. Sprouty genes prevent excessive FGF signalling in multiple cell types throughout development of the cerebellum. Development 138, 2957–2968 (2011).

13. Mahoney Rogers, A.A., Zhang, J. & Shim, K. Sprouty1 and Sprouty2 limit both the size of the otic placode and hindbrain Wnt8a by antagonizing FGF signaling. Developmental biology 353, 94–104 (2011).

14. Kuracha, M. R., Siefker, E., Licht, J. D. & Govindarajan, V. Spry1 and Spry2 are necessary for eyelid closure. Developmental biology 383, 227–38 (2013).

15. Charles, C. et al. Regulation of tooth number by fine-tuning levels of receptor-tyrosine kinase signaling. Development 138, 4063–4073 (2011).

16. Ching, S. T., Cunha, G. R., Baskin, L. S., Basson, M. A. & Klein, O. D. Coordinated activity of Spry1 and Spry2 is required for normal development of the external genitalia. Dev. Biol. 386, 1–11 (2014).

17. Hoshi, M., Batourina, E., Mendelsohn, C. & Jain, S. Novel mechanisms of early upper and lower urinary tract patterning regulated by RetY1015 docking tyrosine in mice. Development 139, 2405–2415 (2012).

18. Alarid, E. T. et al. Keratinocyte growth factor functions in epithelial induction during seminal vesicle development. Proceedings of the National Academy of Sciences 91, 1074–1078 (1994).

19. Thomson, A. A. & Cunha, G. R. Prostatic growth and development are regulated by FGF10. Development 126, 3693–3701 (1999).

20. Zhao, F. et al. Elimination of the male reproductive tract in the female embryo is promoted by COUP-TFII in mice. Science 357, 717–720 (2017).

21. Hanafusa, H., Torii, S., Yasunaga, T. & Nishida, E. Sprouty1 and Sprouty2 provide a control mechanism for the Ras/MAPK signalling pathway. Nat Cell Biol 4, 850–858 (2002).

22. Fong, C. W. et al. Tyrosine phosphorylation of Sprouty2 enhances its interaction with c-Cbl and is crucial for its function. The Journal of biological chemistry 278, 33456–64 (2003).

23. Hall, A. B. et al. hSpry2 Is Targeted to the Ubiquitin-Dependent Proteasome Pathway by c-Cbl. Current Biology vol. 13 308–314 (2003).

24. Rubin, C. et al. Sprouty Fine-Tunes EGF Signaling through Interlinked Positive and Negative Feedback Loops. Current Biology 13, 297–307 (2003).

25. Shim, K., Minowada, G., Coling, D. E. & Martin, G. R. Sprouty2, a mouse deafness gene, regulates cell fate decisions in the auditory sensory epithelium by antagonizing FGF signaling. Dev Cell 8, 553–564 (2005).

26. Vaquero, M. et al. Sprouty1 haploinsufficiency accelerates pheochromocytoma development in Pten+/-mice. Endocrine-related cancer (2016) doi:10.1530/ERC-15-0585.

27. Kuslak, S. L., Thielen, J. L. & Marker, P. C. The mouse seminal vesicle shape mutation is allelic with Fgfr2. Development 134, 557–565 (2007).

28. Okazawa, M. et al. Region-specific regulation of cell proliferation by FGF receptor signaling during the Wolffian duct development. Dev Biol 400, 139–147 (2015).

29. Ornitz, D. M. & Itoh, N. New developments in the biology of fibroblast growth factors. WIREs Mechanisms of Disease n/a, e1549.

30. Guo, L., Degenstein, L. & Fuchs, E. Keratinocyte growth factor is required for hair development but not for wound healing. Genes Dev. 10, 165–175 (1996).

31. Donjacour, A. A., Thomson, A. A. & Cunha, G. R. FGF-10 plays an essential role in the growth of the fetal prostate. Dev Biol 261, 39–54 (2003).

32. Muñoz-Espín, D. et al. Programmed Cell Senescence during Mammalian Embryonic Development. Cell 155, 1104–1118 (2013).

33. Storer, M. et al. Senescence Is a Developmental Mechanism that Contributes to Embryonic Growth and Patterning. Cell 155, 1119–1130 (2013).

34. Muñoz-Espín, D. et al. Programmed Cell Senescence during Mammalian Embryonic Development. Cell 155, 1104–1118 (2013).

35. Murata, T. et al. β-catenin C429S mice exhibit sterility consequent to spatiotemporally sustained Wnt signalling in the internal genitalia. Scientific Reports 4, 6959 (2015).

36. Klein, O. D. et al. Sprouty genes control diastema tooth development via bidirectional antagonism of epithelial-mesenchymal FGF signaling. Dev Cell 11, 181–190 (2006).

37. Chao, C.-M. et al. Fgf10 deficiency is causative for lethality in a mouse model of bronchopulmonary dysplasia. J Pathol 241, 91–103 (2017).

38. Messal, H. A. et al. Antigen retrieval and clearing for whole-organ immunofluorescence by FLASH. Nat Protoc 16, 239–262 (2021).

