## Supplemental Figure and Legends for "A dominant negative mutation uncovers cooperative control of caudal Wolffian Duct development by Sprouty genes"

Supplemental Figure 1. Altés et al.

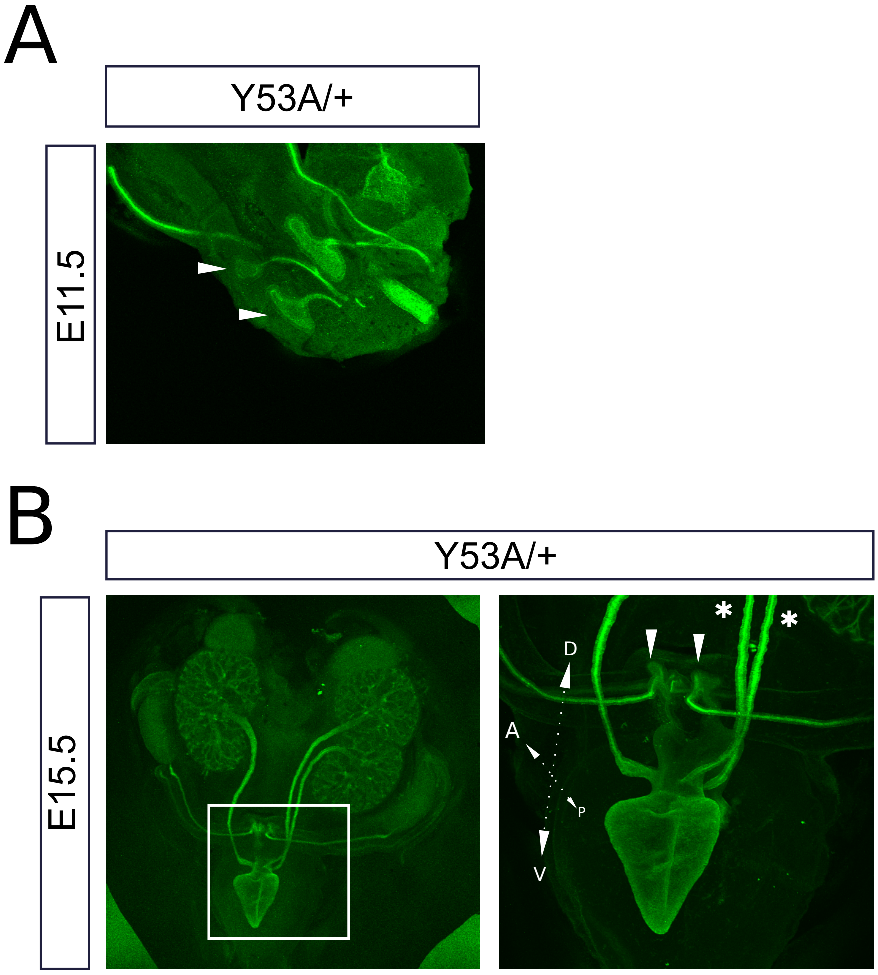

Supplemental Figure 2. Altes et al.

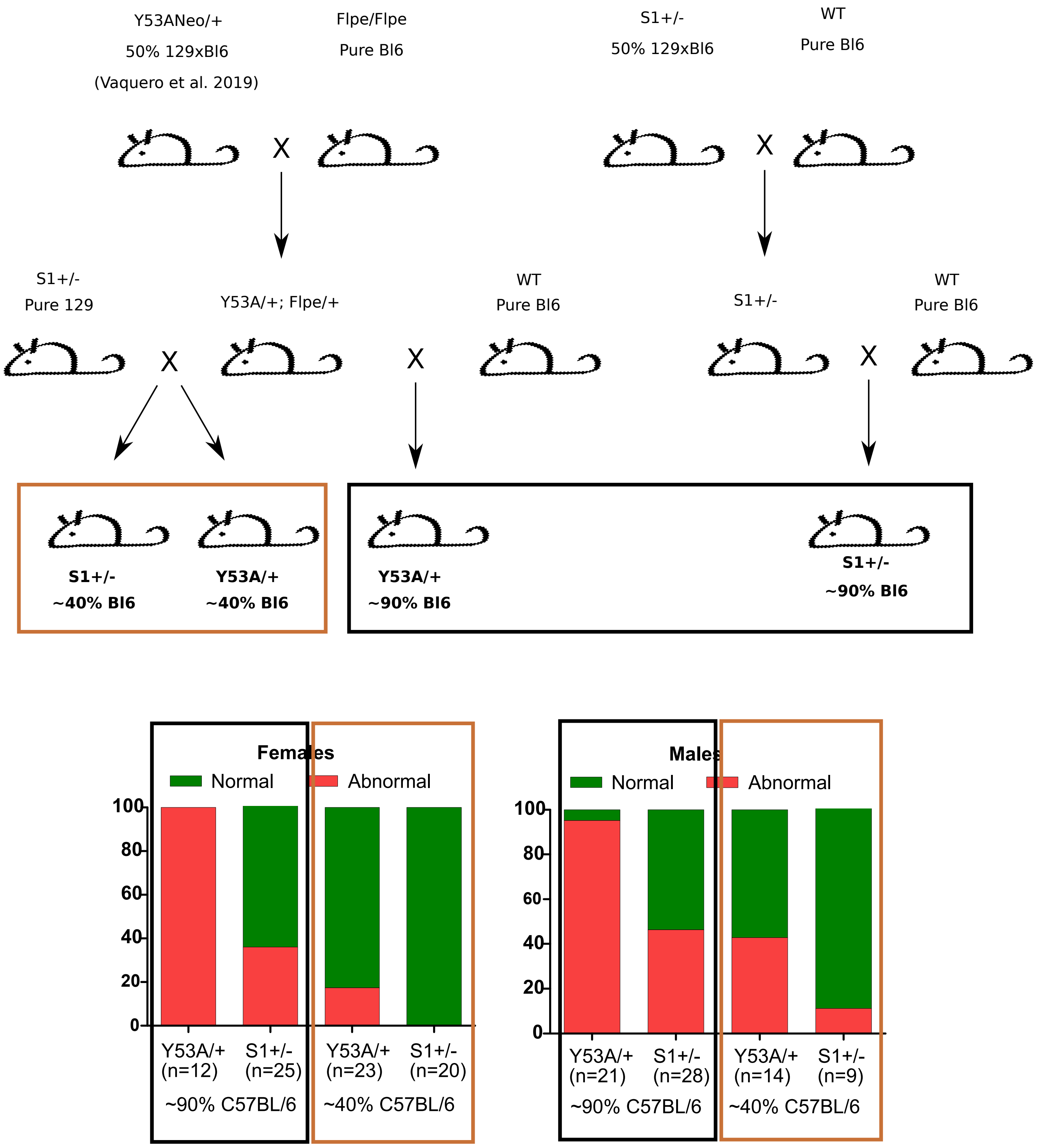

Supplemental Figure 3. Altes et al.

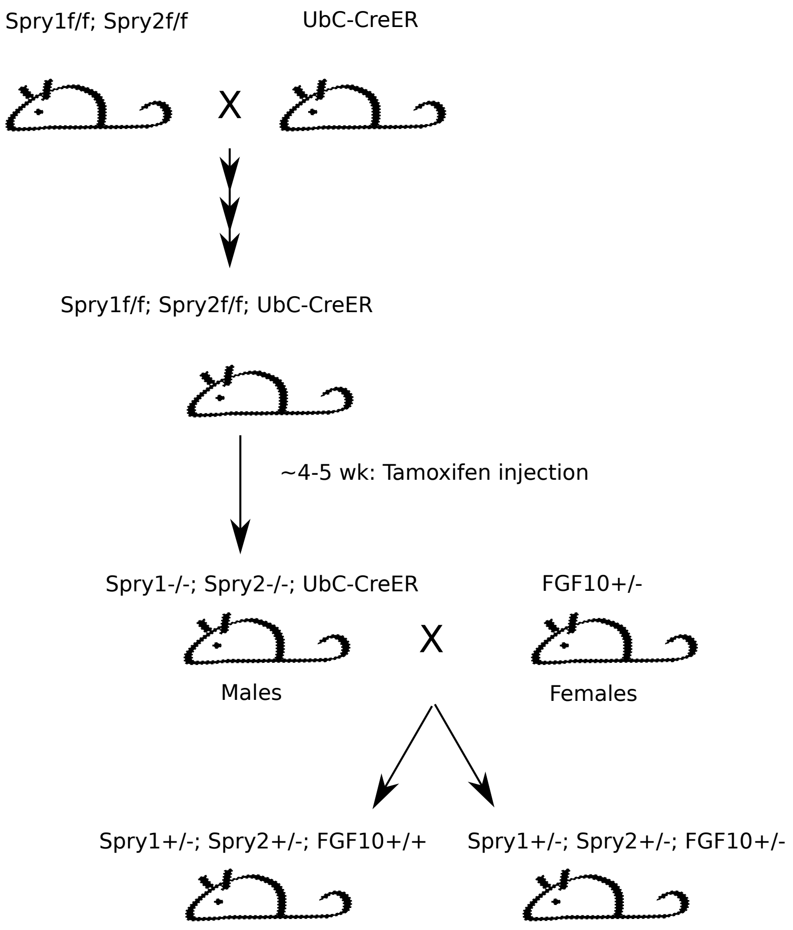

**Supplemental video 1.** Cytokeratin staining of E14.5 wild type lower urinary tract showing complete separation of WD and ureters.

**Supplemental video 2.** Cytokeratin staining of E14.5 Spry1Y$^{53A/+}$ lower urinary tract showing proper ureter maturation.

**Supplemental video 3.** Cytokeratin staining of internal genitalia from a newborn Spry1Y$^{53A/+}$ female mouse showing abnormal retention of the WD along the length of the vagina.

**Supplemental Figure 1.** A small fraction of Spry1Y53A/+ mice display unilateral ectopic UBs generating duplex ureters that are properly segregated from WDs. Whole-mount cytokeratin staining of (A) E11.5 (arrowheads point to duplicated UBs) and (B) E15.5 Spry1Y53A/+ embryos. Asterisks denote duplex ureter and arrowheads show correctly inserted WDs. A→P, and D→V: antero-posterior and dorso-ventral axes, respectively.

**Supplemental Figure 2.** Breeding schemes used to generate Spry1Y53A/+ and Spry1+/- mice in ~50% and ~90% C57BL/6 genetic backgrounds

**Supplemental Figure 3.**.Breeding schemes used to generate Spry1+/-; Spry2+/- mice lacking or not one copy of Fgf10.

**Supplemental Table 1.** Quantification of expression of RTKs in pooled seminal vesicles from 4 newborn wild type animals. RNASeq data are expressed as reads per kilobase (RPK) and transcripts per million (TPM). Fifth column, the position of each gene among the 20150 genes whose expression was detected. Last column indicates the percentile of each gene.

**Supplemental table 2.** Quantification of expression of FGFs, their receptors and FGF binding proteins in pooled seminal vesicles from 4 newborn wild type animals. RNASeq data are expressed as reads per kilobase (RPK) and transcripts per million (TPM).
